# Functional Connectivity Between Somatosensory and Motor Brain Areas Predicts Individual Differences in Motor Learning by Observing

**DOI:** 10.1101/110924

**Authors:** Heather R. McGregor, Paul L. Gribble

## Abstract

Action observation can facilitate the acquisition of novel motor skills, however, there is considerable individual variability in the extent to which observation promotes motor learning. Here we tested the hypothesis that individual differences in brain function or structure can predict subsequent observation-related gains in motor learning. Subjects underwent an anatomical MRI scan and resting-state fMRI scans to assess pre-observation grey matter volume and pre-observation resting-state functional connectivity (FC), respectively. On the following day, subjects observed a video of a tutor adapting her reaches to a novel force field. After observation, subjects performed reaches in a force field as a behavioral assessment of gains in motor learning resulting from observation. We found that individual differences in resting-state FC, but not grey matter volume, predicted post-observation gains in motor learning. Pre-observation resting-state FC between left S1 and bilateral PMd, M1, S1 and left SPL was positively correlated with behavioral measures of post-observation motor learning. Sensory-motor resting-state FC can thus predict the extent to which observation will promote subsequent motor learning.

**New & Noteworthy:** We show that individual differences in pre-observation brain function can predict subsequent observation-related gains in motor learning. Pre-observation resting-state functional connectivity within a sensory-motor network may be used as a biomarker for the extent to which observation promotes motor learning. This kind of information may be useful if observation is to be used as a way to boost neuroplasticity and sensory motor recovery for patients undergoing rehabilitation for diseases that impair movement such as stroke.

## Introduction

Recent work has shown that action observation can promote motor learning. For example, individuals can learn how to reach in novel robot-imposed force field (FF) environments by observing the movements of a tutor (Mattar and Gribble, 2005). Subjects observed a video of a tutor adapting his reaches to a novel robot-imposed FF applied. Subjects who later performed reaches in the same FF showed a benefit, performing better (straighter) reaches compared to control subjects who did not observe a tutor. Subjects who later performed reaches in the opposite FF performed worse (more curved) reaches than subjects who did not observe. While these results demonstrate that FFs can be partially learned from observation, there is considerable inter-individual variability in the extent to which observation promotes motor learning. Little is known about why this may be. Some individuals may be more predisposed to learning from observation than others, whether from birth, from experience-dependent plasticity, or a combination of these or other individual differences. Here we test the idea that inter-individual differences in brain function or structure underlie the extent to which observation promotes subsequent motor learning.

In a recent review article, Zatorre (2013) discusses findings showing how structural and functional neural connectivity patterns predict individual differences in musical training and speech learning. Other studies have shown similar predictability for a wide array of cognitive abilities including executive function (Barnes et al., 2014; Reineberg et al., 2015), reading (Koyama et al., 2011; Wang et al., 2013), second language acquisition (Chai et al., 2016), visual perceptual discrimination (Baldassarre et al., 2012) and memory recall (King et al., 2015). In the motor domain, Tomassini et al. (2011) demonstrated that individual differences in both functional and structural magnetic resonance imaging (MRI) measures correlate with the acquisition of a novel visuomotor tracking skill through active movement training. Task-based functional activation levels in a network involving prefrontal, premotor, and parietal cortices, as well as basal ganglia and the cerebellum were associated with behavioral measures of active motor learning. Structural differences within the premotor cortex, higher order visual areas, and the cerebellum were also positively correlated with learning abilities (Tomassini et al., 2011). Similarly, using dense-array electroencephalography (EEG), Wu et al. (2014) showed that resting-state functional connectivity (FC) between premotor, primary motor and parietal cortices predicts individual differences in the subsequent learning of a visuomotor tracking task. Together, these studies suggest that functional and structural variations in motor learning-related brain networks can, in part, explain individual differences in the ability to learn novel motor tasks through active movement practice. The results of these studies raise the possibility that individual differences in brain structure or function may also be predictive of motor learning by observing.

Here we tested the hypothesis that individual differences in brain function or structure can predict the extent to which individuals will learn to perform a novel sensory-motor task (FF reaching) from observation. Based on our previous work (McGregor and Gribble, 2015; McGregor et al., 2016), we expected that individual differences in brain function and structure within visual and sensory-motor brain networks would be predictive of motor learning by observing. On day 1, subjects performed baseline (no FF) reaches using a robotic arm and then underwent pre-observation anatomical and resting-state functional magnetic resonance imaging (fMRI) scans. Twenty-four hours later, subjects in a learning group observed a video of a tutor learning to reach in a novel FF. Subjects in a control group observed a video of a tutor performing reaches in an unlearnable FF. Following observation, all subjects performed reaches in a FF as a behavioral assessment of motor learning by observing. We found that, for the learning group, pre-observation (day 1) resting-state FC between bilateral dorsal premotor cortex (PMd), primary motor cortex (M1), primary somatosensory cortex (S1) and left superior parietal lobule (SPL) was reliably correlated with behavioral scores of motor learning by observing acquired on day 2. No such correlation between pre-observation FC and motor learning by observing scores was found for the control group. Moreover, we found that individual differences in grey matter volume could not predict subsequent motor learning by observing. Pre-observation sensory-motor resting-state FC can thus explain part of the between-subject variation in motor learning by observing.

## Materials and Methods

### Subjects

Thirty healthy subjects participated in this study. Fifteen subjects were assigned to a learning group (6 males, mean age 22.87 ± 1.02 (SE) years) and 15 were assigned to a control group (6 males, mean age 22.53 ± 0.86 (SE) years). All subjects were right handed, had normal or corrected-to-normal vision, were naïve to force fields, and reported no neurological or musculoskeletal disorders. Subjects provided written informed consent prior to participating. All experimental procedures were approved by the Research Ethics Board at the University of Western Ontario.

### Apparatus

Subjects were seated in front of a custom tabletop and grasped the handle of a two degree of freedom robotic arm (IMT2, Interactive Motion Technologies) with the right hand (see Figure 1). The chair height was adjusted such that the subject’s upper arm was abducted approximately 90^o^ from the trunk. An air sled was secured beneath the subject’s right arm to support the arm against gravity. A semi-silvered mirror, mounted horizontally just above the robotic arm, occluded the subject’s vision of his or her own arm and the robotic arm. During the reaching task, a liquid crystal display television (LCD TV) projected visual feedback onto the semi-silvered mirror. Visual feedback included a start position (20-mm blue circle), a single target (20-mm white circle), and a cursor representing hand position (12-mm pink circle).

**Figure 1:**
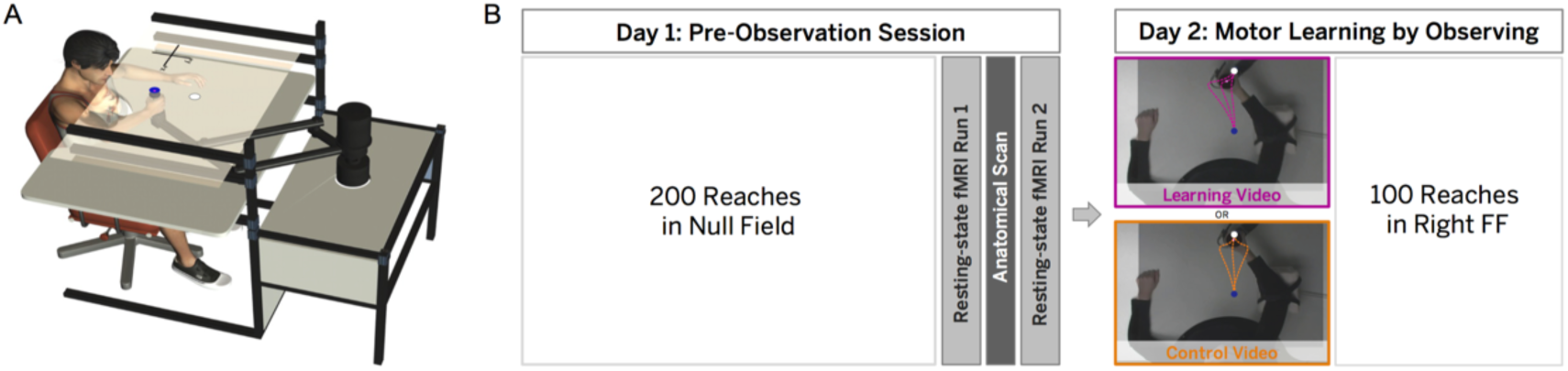
Apparatus and Experimental Design. **A.** Subjects were seated in front of an InMotion2 robotic arm and performed the reaching task in a horizontal plane using the right arm. **B.** On day 1, all subjects performed reaches in a null field (no force applied by the robot). Subjects then underwent a pre-observation MRI scan session. The scan session consisted of 2 resting-state runs separated by an anatomical scan, followed by 2 functional localizer tasks. On day 2, subjects in the learning group (n = 15) observed a learning video showing a tutor adapting her reaches to a left FF. A control group (n = 15) observed a control video showing a tutor performing curved reaches in an unlearnable (randomly-varying) FF. Finally, all subjects performed reaches in a right FF as a behavioral test of motor learning by observing. FF, force field.

The reaching task involved guiding the handle of the robotic arm from the start position to the target, which was located 15 cm in front of the start position. Subjects were instructed to move as straight as possible. At the end of each reach, the target changed color to provide feedback about movement time: the target disappeared if the movement time was within the desired range (450-550 ms duration), turned red if the movement was too fast (< 450 ms) or turned green if the movement was too slow (> 550 ms). Following each reach, the robotic arm returned the subject’s hand to the start position.

The robot applied a velocity-dependent force field during the reaching task according to Equation 1:

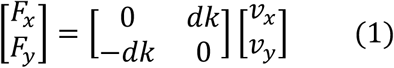

in which *x* and *y* are lateral and sagittal directions, *F_x_* and *F_y_* are the applied robot forces,*v_x_* and *v_y_* are hand velocities, *k*=14 Ns/m, and *d*=0 (null field), +1 (right FF) or −1 (left FF).

### Reaching Video Stimuli

Each video showed a top-down view of a tutor performing the reaching task described above using her right arm. The tutors in the videos were naïve to force fields. The learning video consisted of a series of 30-second clips showing a tutor adapting her reaches to a leftward force field (left FF). These clips showed the gradual progression from curved to straight movements that is indicative of motor learning. The control video consisted of a series of 30-second clips showing a tutor performing reaches in an unlearnable FF in which the direction of the FF varied randomly from trial to trial (left FF, right FF, or null field). These clips showed the tutor performing both high and low curvature movements, but lacked the progressive decrease in movement curvature depicted in the learning video. Therefore, the control video included similar movements to those shown in the learning video, but did not depict learning. The videos showed 200 reaches each and were 15 minutes in duration (including regular breaks). Video screenshots are shown in Figures 1B and 2A. Note that the dashed trajectories and superimposed labels have been included for demonstrative purposes here and were not shown to subjects in the experiment.

**Figure 2:**
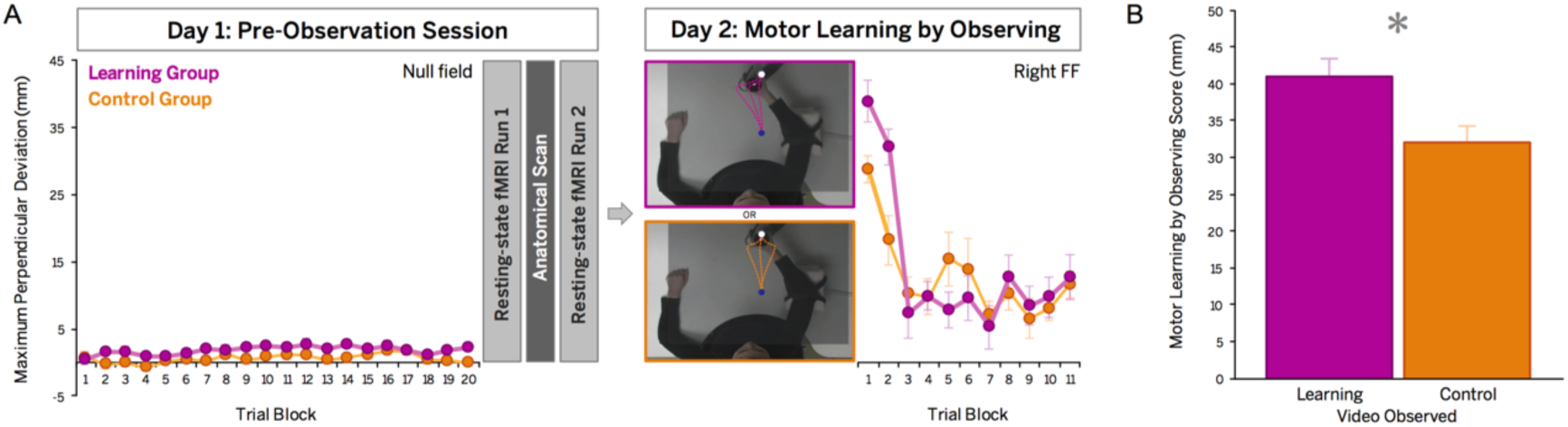
Behavioral results. **A.** Experimental design showing the average PD of reaches for each group across trials in the null field on day 1 and in the right FF on day 2. Behavioral data from the learning and control group are shown in magenta and orange, respectively. Data are shown as 10-trial blocks except for the first 2 blocks in the right FF, which are shown as 5-trial blocks. Error bars represent SEM. **B.** Motor learning by observing scores for the learning group (magenta) and control group (orange), reflecting initial PD in the right FF relative to baseline PD in the null field. Error bars represent SEM. FF, force field; PD, perpendicular deviation.

### Experimental Design

The experimental design is shown in Figure 1B. All subjects (n=30) participated in three sessions. For each subject, the sessions were held at the same time on three consecutive days. On day 0, subjects were familiarized with the reaching task by performing 50 practice movements in a null field (no force applied by the robot). On day 1, subjects performed 200 baseline reaches in the null field and then walked to the imaging facility for a fMRI scan session. The scan session, described in detail below, began approximately 20 minutes following the completion of the reaching task and lasted 1 hour. Data collected during the day 1 scan session were used to estimate pre-observation resting-state FC involving 10 visual and sensory-motor brain areas (see ROIs below) and to estimate whole-brain grey matter volume. On day 2, subjects performed the observational motor learning task. Subjects watched either the learning video or the control video while seated in front of the robotic arm. The video was played on the LCD TV positioned above the robotic arm and was projected onto the semi-silvered mirror surface. To ensure subjects paid attention during the video, we instructed them to count the number of correctly-timed reaches in the video (indicated by the target disappearing upon the completion of a reach) and to report the final tally to the experimenter following the video. Reported tallies were analyzed to verify that subjects attended to the video, but these data were not incorporated into the behavioral or neuroimaging analyses. Note that subjects were not told to pay attention to any particular part of the movement trajectory or arm, nor were they told that the robot would be applying forces to the arm. Approximately 80 minutes after video observation, we assessed motor learning by observing by having subjects perform 100 reaches while the robotic arm applied a rightward FF (right FF).

During the 80 minutes between video observation and the motor learning test on day 2, both groups underwent a second fMRI scan session identical to the day 1 fMRI scan session. Data from the second fMRI scan session were not used in any of the analyses presented here since the main objective of the current study was predicting motor learning by observing based on pre-observation (day 1) neuroimaging data. Using this same dataset, we have previously examined changes in resting-state FC from pre-observation (day 1 scan) to post-observation (day 2 scan). See McGregor and Gribble (2015) for details of FC changes from day 1 to day 2, and how they relate to observation-related gains in motor learning.

We assessed motor learning behaviorally by having subjects perform reaches in a right FF, which was the opposite FF to what was depicted in the learning video. The more subjects learned about the observed left FF, the worse their performance would be in the right FF. The idea is that, during observation, subjects learn about the compensatory pattern of muscle forces (i.e., rightward compensation) that is required to counteract the left FF. Subjects use this learned pattern of muscle forces when they subsequently perform reaches, resulting in after-effects (e.g., Shadmehr and Mussa-Ivaldi, 1994). As is the case in this study, after-effects are especially large if the FF is changed such that it is the opposite of the learned environment. This is because the subject compensates rightward (persistence of the learned pattern of muscle forces) and the robotic arm also pushes the hand to the right. Therefore, we expected that those subjects who better learned about the observed left FF would perform more highly curved reaches when first exposed to the right FF (Cothros et al., 2006; Brown et al., 2009; McGregor and Gribble, 2015; McGregor et al., 2016). We chose to use this interference paradigm to assess motor learning by observing because it tends to be a more sensitive measure compared to testing subjects in the same FF that they observed.

### Imaging Procedure

Neuroimaging data were acquired by a 3-Tesla Siemens Magnetom Tim Trio imaging system using a 32-channel head coil. The fMRI scan session lasted 1 hour. The scan session began with two 8-minute resting-state runs during which subjects were instructed to relax with their eyes closed. The resting-state runs were separated by a 5-minute anatomical scan during which subjects were instructed to fixate their gaze on a crosshair projected onto a screen. Subjects then performed two 6-minute functional localizer tasks: an action observation network localizer task and a motor localizer task. We selected 10 *a priori* regions of interest (ROIs) known to be involved in action observation and/or motor learning (see below). The two localizer tasks allowed us to determine the coordinates of each ROI for use in the functional connectivity analysis described below.

For the the action observation network localizer task, subjects viewed intact and scrambled video clips of a tutor performing reaches while holding the robotic arm (ten 36-s interleaved blocks in total). Intact video clips showed a top-down view of a tutor performing straight reaching movements in a null field (no forces applied by the robot). For the baseline condition, subjects viewed scrambled versions of these video clips in which only the start and target positions remained in their original locations. Scrambling the videos allowed us to preserve the low-level motion features such as movement direction and velocity while removing such movement features as shoulder and elbow joint rotations and the hand path (Malfait et al., 2010). During the action observation network localizer task, subjects were instructed to count the number of correctly-timed movements the tutor performed and to report the final tally to the experimenter at the end of the video. This was done to verify that subjects attended to the video. Reported tallies were not incorporated into the behavioral or neuroimaging analyses.

For the motor localizer task, subjects performed interleaved blocks of arm movement and rest (ten 36-s blocks in total). During movement blocks, subjects slowly moved their right forearm along the frontal plane in a cyclic manner (90^o^ elbow flexion). Color-coded visual cues were used to pace movements at a frequency of 0.1 Hz.

### Image Acquisition

Whole-brain functional data were acquired with a T2-weighted EPI sequence (TR = 3,000 ms, TE = 30 ms, 90^o^ flip angle, 3-mm isotropic voxels, 80×80×50 matrix, iPAT acceleration factor = 2). T1-weighted anatomical images were collected with a MPRAGE sequence (TR = 2,300 ms, TE = 2.98 ms, 9^o^ flip angle, 1-mm isotropic voxels, 192×240×256 matrix). For each subject, a field map was acquired at the beginning of the scan session using a gradient echo sequence (TR = 531 ms, TE = 4.92 ms/7.38 ms, 60^o^ flip angle, 3-mm isotropic voxels, 80×80×50 matrix).

### Behavioral Data Analysis

During the reaching task, the position and velocity of the robotic handle were sampled at 600 Hz and stored for offline analysis. Positional data were low-pass filtered offline at 40 Hz. The start and end of each trial were defined using a threshold of 5% of the peak tangential velocity of the hand. Movement curvature was quantified for each trial as the maximum perpendicular deviation (PD) of the hand from a straight line connecting the start and target locations (Mattar and Gribble, 2005).

We calculated a behavioral motor learning by observing score for each subject. The Motor learning by observing scores were calculated as the mean PD of the first 3 reaches in the right FF minus the mean PD of the last 50 reaches in the baseline null field. This approach allowed us to examine the extent to which observing the left FF interfered with subjects' initial performance in the right FF compared to control subjects who did not observe the tutor undergoing learning. As in our previous work (Cothros et al., 2006; Brown et al., 2009; McGregor et al., 2016), we expected that motor learning by observing would primarily affect initial performance in the right FF, after which motor learning through active movement in the right FF would occur for both groups.

### Functional Connectivity Analysis

We carried out a whole-brain seed-based correlation analysis to examine if inter-subject differences in resting-state FC on day 1 could predict the amount of motor learning by observing that subjects would achieve on the following day. Neuroimaging data analyses were performed using FSL version 5.04 (FMRIB's Software Library, https://www.fmrib.ox.ac.uk/fsl). Image preprocessing steps for the functional connectivity analysis included the removal of the first 2 volumes in each functional run, slice-timing correction, motion correction, spatial smoothing using a 6-mm kernel, and high-pass temporal filtering (100 s). Field map distortion correction and affine coregistration of functional and anatomical images were performed using boundary-based registration (BBR) in FLIRT. Subjects' images were registered to MNI standard space (MNI's 152-brain T1 template, 2-mm isotropic voxel size) using a 12-DOF affine registration.

Following preprocessing, each resting-state run was bandpass filtered between 0.01 Hz and 0.1 Hz (Biswal et al., 1995; Damoiseaux et al., 2006). Mean-based intensity normalization was performed (mean value of 10,000) to remove global intensity differences between runs (Damoiseaux et al., 2006). We then carried out our seed-based correlation analysis using FILM (FMRIB's Improved General Linear Model).

We selected 10 *a priori* regions of interest (ROIs) known to be involved in action observation and/or motor learning. ROIs included left supplementary motor area (SMA), dorsal premotor cortex (PMd), ventral premotor cortex (PMv), primary motor cortex (M1), primary somatosensory cortex (S1), visual area V5/MT, superior parietal lobule (SPL), inferior parietal lobule (IPL), putamen, and right cerebellum. We determined the coordinates of each ROI based on the results of the block-design analyses of the action observation network localizer task and the motor localizer task. For each localizer, the task-induced response was assessed with a per-subject GLM. Data from all 30 subjects were then included in a mixed-effects analysis (Z > 2.3, p < 0.05, cluster-based thresholding) for each localizer. These analyses yielded Z-score maps showing areas of the brain that were activated (on average across all 30 subjects) during arm movement or action observation, which we used to determine the coordinates of our ROIs. For each of our 10 ROIs, we found the peak activated voxel within that brain area and centered the ROI on that voxel. Each ROI consisted of all voxels within a 6-mm radius of the activation peak. Table 1 shows the coordinates of the activation peaks on which each ROI was centered.

**Table 1:**
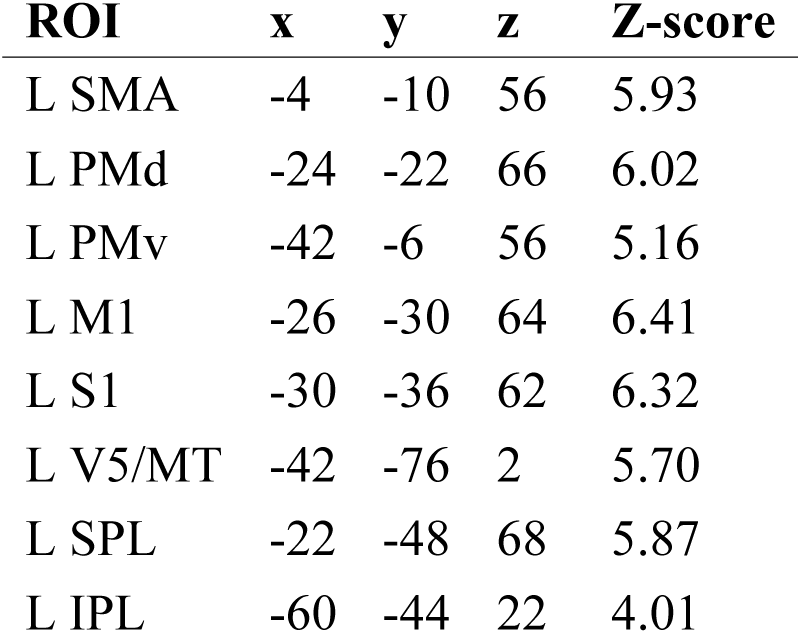
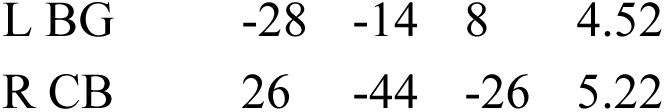
Region of Interest (ROI) coordinates used in functional connectivity analyses for both the learning and control groups. The ROI coordinates were determined on the basis of a block-design analysis of the action observation network localizer task and motor localizer task. The seed coordinates were chosen as the peak activated voxel within each of the 10 a priori selected brain regions listed in this table. L, left; R, right; SMA, supplementary motor area; PMd, dorsal premotor cortex; PMv, ventral premotor cortex; M1, primary motor cortex; S1, primary somatosensory cortex; V5/MT, middle temporal visual area; SPL, superior parietal lobule; IPL, inferior parietal lobule; BG, putamen; CB, cerebellum. ROI locations are given in the MNI coordinate frame.

| ROI | x | y | z | Z-score |
| --- | --- | --- | --- | --- |
| L SMA | -4 | -10 | 56 | 5.93 |
| L PMd | -24 | -22 | 66 | 6.02 |
| L PMv | -42 | -6 | 56 | 5.16 |
| L M1 | -26 | -30 | 64 | 6.41 |
| L S1 | -30 | -36 | 62 | 6.32 |
| L V5/MT | -42 | -76 | 2 | 5.70 |
| L SPL | -22 | -48 | 68 | 5.87 |
| L IPL | -60 | -44 | 22 | 4.01 |
| L BG | -28 | -14 | 8 | 4.52 |
| R CB | 26 | -44 | -26 | 5.22 |

We then carried out a functional connectivity analysis to estimate FC between each ROI and the rest of the brain on day 1. For each ROI, we carried out a subject-level analysis on each resting-state run in which the mean time series of the ROI was used as the predictor of interest. Nuisance regressors included the temporal derivative of the mean ROI time series, 6 rigid body motion parameters obtained from motion correction, mean global signal, mean white matter signal and mean CSF signal. The results of the subject-level analyses were then entered into a mixed-effects group-level analysis for each of the ROIs. A separate mixed-effects analysis was carried out for each group. In the group-level analysis, we also included a nuisance regressor modeling inter-subject differences in baseline movement curvature in the null field. This nuisance regressor consisted of each subject’s average PD of the last 50 reaches in the null field. This was done because subjects had performed 200 reaches in the null field prior to the fMRI scan session on day 1. Even though the robot did not apply forces to the hand during null field reaches, subjects likely underwent some degree of motor learning as they learned the inertial properties of the robotic arm. We included the nuisance regressor modeling subjects' behavioral performance in the null field to account for variability in pre-observation resting-state FC that could be explained by differences in subjects' movement curvature at baseline (in the null field condition). Group-level functional connectivity analyses were performed using both of the subjects' resting-state runs together as well as separately (see Results).

Group-level analysis results were thresholded based on Gaussian random field theory using a maximum height thresholding (Z >5.3) with a corrected significance level of p = 0.005 (voxelwise thresholding, corrected for familywise error). We applied a Bonferroni correction for the number of ROIs used; therefore, our corrected significance threshold of p = 0.005 reflects p = 0.05/10 ROIs. These analyses resulted in 10 Z score maps per group (one per ROI) showing areas that, on average, exhibited FC with the seed region across subjects.

For each of the 10 resulting Z score maps, FC was computed for each subject in the group as the temporal correlation (Fisher Z-transformed correlation coefficient) between the ROI time series and the average time series across all clusters in the identified network. This allowed us to estimate each subject’s day 1 FC between the ROI and all of the clusters in each of the identified networks. At the group-level, we computed the correlation (across subjects) between day 1 FC values and day 2 motor learning by observing scores for each of the identified networks. This was done to assess if individual differences in day 1 FC among brain areas in any of the identified networks was related to performance during the behavioral test of motor learning by observing on day 2. We again applied a Bonferroni correction for the number of ROIs used; therefore, we considered statistically significant only those correlations between day 1 FC and motor learning by observing scores for which p<0.005 (i.e., p=0.05/10 ROIs).

### Voxel-Based Morphometry Analysis

We carried out a whole-brain voxel-based morphometry (VBM) analysis to test for inter-subject differences in grey matter volume across the whole brain (measured on day 1) that could predict motor learning by observing scores on day 2. This analysis was carried out on the T1-weighted images using FSL-VBM v1.1. First, each subject’s anatomical image was brain-extracted, grey-matter segmented, and transformed to MNI space using a nonlinear registration. The resulting anatomical images were then averaged and flipped along the x-axis to generate a left-right symmetric, study-specific template. Each subject’s grey matter-segmented anatomical image was registered to the study-specific template and smoothed using a 3-mm Gaussian kernel. The VBM analysis was carried out using a voxelwise GLM model. The predictor of interest modeled the subjects' motor learning by observing scores (demeaned). Two nuisance regressors were also included in the GLM; one modeled the grey matter grand mean across all subjects and the second modeled each subject’s unnormalized total brain volume. Each subject’s total brain volume was estimated prior to standard space normalization using FSL's SIENAX tool. The voxelwise GLM model was applied using non-parametric permutation (50,000 iterations) to correct for multiple comparisons with a significance threshold of p=0.05.

## Results

### Behavioral Results

Figure 2A shows the behavioral data from the learning and control groups. It can be seen that, on day 1, reaches are straight in the baseline null field condition for both groups. Following video observation on day 2, we assessed motor learning by observing by instructing subjects to perform straight reaches while the robotic arm applied a right FF (the opposite FF to what had been observed in the learning video). The more subjects learned about the observed left FF, the worse their performance would be during their initial performance in the right FF. Indeed, we found that subjects who observed the tutor adapting to a left FF in the learning video exhibited greater PD during initial reaches in the right FF compared to control subjects who observed the tutor performing curved reaches in an unlearnable FF. As in previous work (Mattar and Gribble, 2005; Cothros et al., 2006; Brown et al., 2009; Williams and Gribble, 2012; Bernardi et al., 2013; McGregor et al., 2016), the effects of observation are most apparent early in the motor learning test (i.e., the first 10 reaches shown as blocks 1 and 2 in Fig 2A) and diminish as subjects in both the learning and control groups adapt to the right FF. Average motor learning by observing scores are shown in Figure 2B. Motor learning by observing scores reflect the PD of the first 3 reaches in the right FF relative to the subject’s baseline PD in the null field. As shown in Figure 2B, subjects who observed the tutor undergoing left FF learning exhibited significantly higher motor learning by observing scores compared to control subjects who observed the tutor performing reaches in an unlearnable FF (*t*(28)=2.58, p < 0.01).

### Functional Connectivity Analysis

We performed a functional connectivity analysis using the resting-state fMRI data acquired on day 1 to test whether individual differences in pre-observation FC could predict motor learning by observing scores on the following day. Of the 10 ROIs used, only the analysis using the left S1 ROI revealed a network in which pre-observation FC was reliably correlated with day 2 motor learning by observing scores for the learning group. As can be seen in Figure 3, day 1 FC between the left S1 ROI and the average FC across clusters in bilateral PMd, bilateral M1, bilateral S1 and left SPL was positively correlated with day 2 motor learning by observing scores (r=0.76, p=0.001) for the learning group. Subjects with greater pre-observation FC among these areas on day 1 went on to achieve higher motor learning by observing scores on the following day. Table 2 shows cluster activation peaks and statistics for the learning group. For the control group, the analysis using the left S1 ROI revealed a qualitatively similar network consisting of bilateral PMd, bilateral M1, bilateral S1 and left SPL. This is expected because subjects in the learning and control groups have had identical experiences as of the day 1 resting-state scan session. However, for the control group, day 1 FC within the identified network was not reliably correlated with day 2 behavioral motor learning by observing scores (r=−0.43, p=0.67; Figure 5).

**Table 2:** The functional connectivity analysis using the ROI in left S1 (see Table 1), revealed a sensory-motor functional network in which pre-observation (day 1) FC predicted day 2 motor learning by observing scores for the learning group. Z score activation peaks, MNI coordinates and anatomical labels of the sensory-motor clusters in the identified functional network are shown here. ROI, region of interest; L, left; R, right; SMA, supplementary motor area; PMd, dorsal premotor cortex; M1, primary motor cortex; S1, primary somatosensory cortex; SPL, superior parietal lobule; FC, functional connectivity.

| **Z-score** | **x** | **y** | **z** | **Label** |
| --- | --- | --- | --- | --- |
| 7.40 | -26 | -40 | 58 | L S1 (BA2) |
| 6.75 | -22 | -20 | 66 | L PMd (BA6) |
| 6.12 | -16 | -50 | 62 | L SPL (BA5L) |
| 5.80 | -34 | -28 | 52 | L M1 (BA4p) |
| 6.97 | 22 | -42 | 60 | R S1 (BA2) |
| 7.10 | 30 | -32 | 56 | R M1 (BA4p) |
| 6.36 | 26 | -16 | 64 | R PMd (BA6) |

Our computed motor learning by observing score took into account the average PD of a subject’s first 3 reaches in the right FF relative to his or her baseline PD in the null field. To assess the sensitivity of the learning group’s correlation between pre-observation FC and motor learning by observing scores, we computed additional motor learning by observing scores to use in our analysis. Additional motor learning by observing scores reflected the average PD of the first 4, 5, 6, 7, 8, 9 or 10 reaches in the right FF minus the average PD of the last 50 reaches in the null field. The learning group's correlation between day 1 FC and motor learning by observing scores remained statistically significant for all of the additional measures.

The GLMs used for the group-level functional connectivity analyses included a nuisance regressor modeling each subject’s baseline PD in the null field during the last 50 trials. This nuisance regressor was included to account for variability in pre-observation resting-state FC that could be explained by inter-subject differences in movement curvature at baseline. Our results were consistent whether the null field nuisance regressor reflected the average PD of the last 3, 5, 10 or 50 null field reaches or the average PD of the first 3, 5, 10 or 50 null field reaches.

It is possible that the correlation between pre-observation FC and the day 2 motor learning by observing scores is due to random chance (e.g., spurious correlations in the BOLD time series) and not due to stable individual differences in functional connectivity. To assess this, we repeated the functional connectivity analysis on each of the two resting-state runs separately. The resting-state runs were independent, separated in time by a 5-minute anatomical scan. Again using the ROI in left S1, we found consistent spatial patterns of pre-observation (day 1) FC between left S1, bilateral PMd, M1, S1 and left SPL for both individual runs (see Figure 4). Moreover, for the learning group, the correlation between pre-observation (day 1) FC and day 2 motor learning by observing scores was statistically significant for both resting-state run 1 (r=0.75, p=0.001) and run 2 (r=0.63, p=0.01). Therefore, when performed on the each of the two independent resting-state runs, our analysis yielded similar results both in terms of the spatial extent of the clusters and the correlations with day 2 motor learning by observing scores. It is therefore unlikely that our main result arises from a spurious correlation. For the control group, there was no statistically significant correlation between pre-observation FC during either run 1 or run 2 and motor learning by observing scores (r=−0.38, p=0.15 and r=−0.03, p=0.91, respectively; Figure 5).

**Figure 3:**
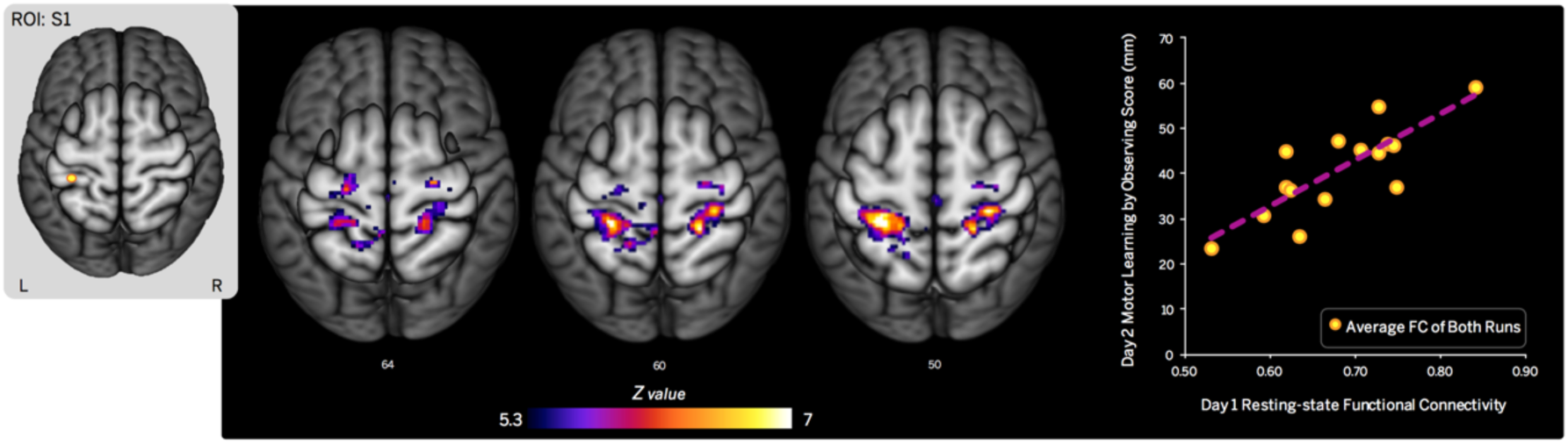
Pre-observation FC predicted motor learning by observing scores for the learning group. This figure shows neuroimaging data from the learning group only. Pre-observation (day 1) resting-state FC between the left S1 ROI (inset at left) and clusters in bilateral PMd, bilateral M1, bilateral S1 and left SPL are shown. FC values reflect the Fisher Z-transformed temporal correlation between the ROI time series and the average time series of all clusters in the identified network for each subject. Across subjects in the learning group, the average day 1 resting-state FC within this network was positively correlated with day 2 motor learning by observing scores. As shown in the scatterplot on the far right, subjects who exhibited stronger resting-state FC within this network on day 1 achieved greater motor learning by observing scores on the following day (r=0.76, p=0.001).

**Figure 4:**
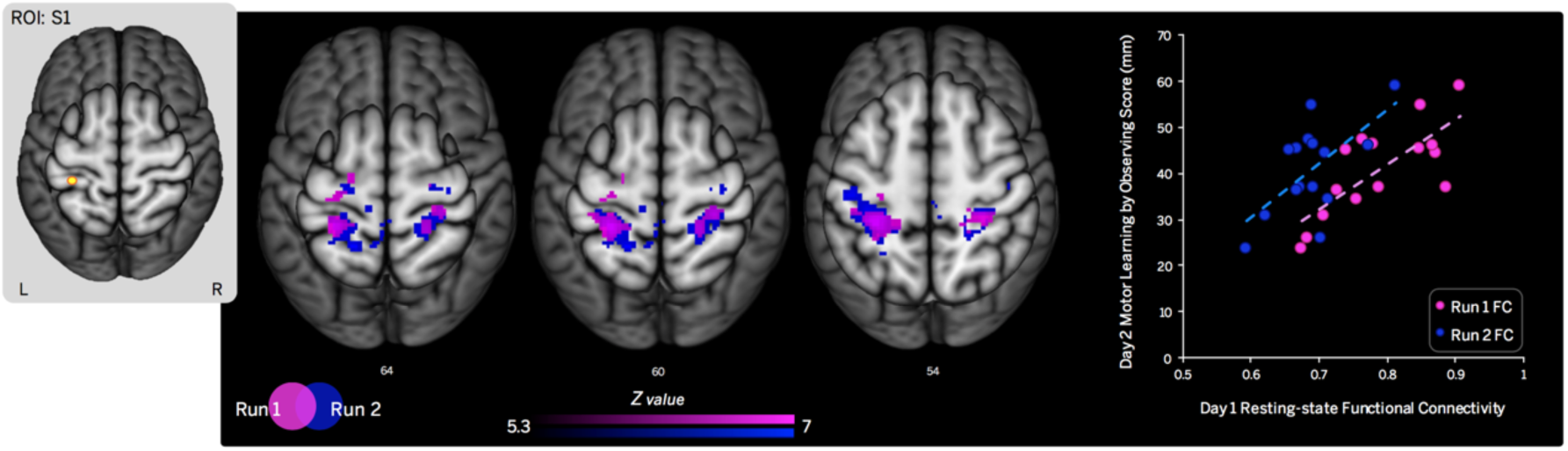
Pre-observation FC in run 1 and run 2 both predicted motor learning by observing scores for the learning group. The figure shows neuroimaging data from the learning group only. Data from resting-state run 1 (shown in pink) and run 2 (shown in blue) were analyzed separately. For each run, the ROI in left S1 (inset at left) exhibited resting-state FC with clusters in bilateral PMd, bilateral M1, bilateral S1 and left SPL. FC values reflect the Fisher Z-transformed temporal correlation between the ROI time series and the average time series of all clusters in the identified network. For each of the runs, pre-observation (day 1) resting-state FC between bilateral PMd, M1, S1 and left SPL was reliably correlated with day 2 motor learning by observing scores across subjects in the learning group. As shown in the scatterplot on the far right, subjects who exhibited stronger FC within the network identified in each run on day 1 achieved greater motor learning by observing scores on day 2.

**Figure 5:**
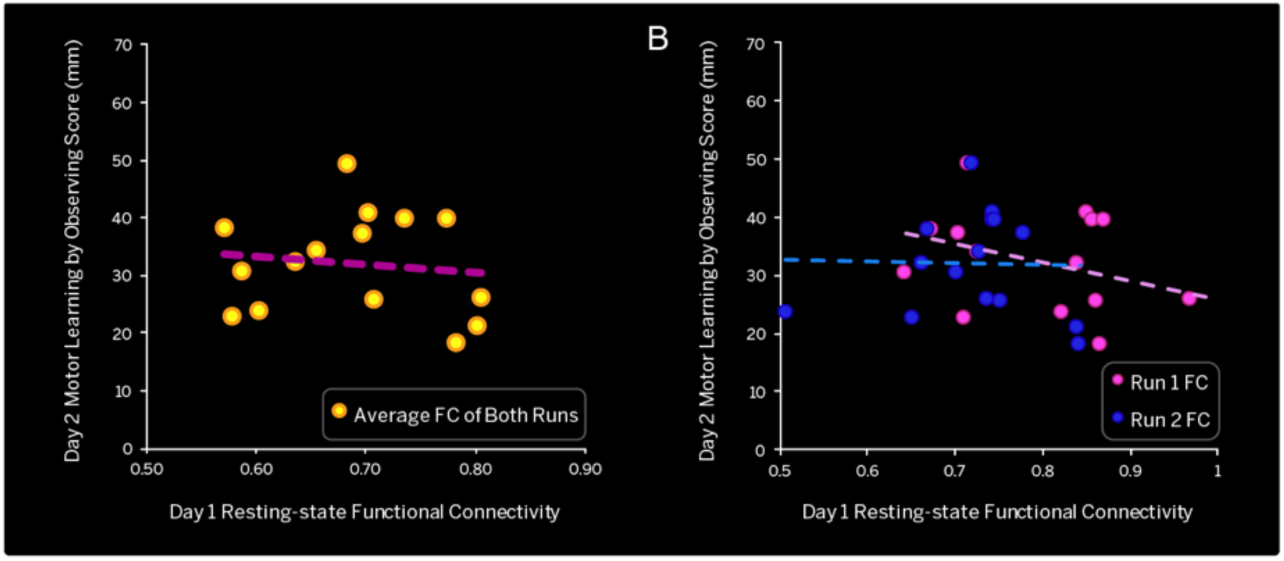
Pre-observation FC did not predict motor learning by observing scores for the control group. As was the case with the learning group, subjects in the control group exhibited pre-observation (day 1) resting-state FC between the left S1 ROI and clusters in bilateral PMd, M1, S1 and left SPL (not shown). FC values reflect the Fisher Z-transformed temporal correlation between the ROI time series and the average time series of all clusters in the identified network. Across subjects in the control group, there was no correlation between day 1 resting-state FC within this network and day 2 motor learning by observing scores. This was the case when both runs were analyzed together (r=−0.38, p=0.15; shown in the scatterplot on the left) as well as when the runs were analyzed separately r=−0.03, p=0.91; shown in the scatterplot on the right).

### Voxel-Based Morphometry Analysis

We carried out a whole-brain VBM analysis on the T1-weighted anatomical images. This was done to test if individual differences in grey matter volume could predict subsequent motor learning by observing scores. This analysis yielded no significant results. We tested the sensitivity of this null result to the chosen statistical threshold. For the learning group, no significant clusters survived statistical thresholding at the group level until the p-value threshold was raised to 0.27, at which level clusters survived in left frontal lobe (−32, 54, 12) and Broca's area (−50, 20, 12). When the p-value threshold was raised further to 0.37, a cluster survived which spanned right PMC (54, −8, 52), M1 (54, −10, 46), S1 (56, −14, 44) and IPL (64, −20, 40). However, since none of these clusters survived an appropriate statistical threshold, these results are not interpretable. In the context of the dataset here, individual differences in grey matter volume could not account for variability in the extent to which observation promotes motor learning.

## Discussion

Here we examined if pre-observation measures of brain function or structure could account for individual differences in the extent to which observation facilitates motor learning. We acquired measures of resting-state FC and grey matter volume using MRI prior to an observational learning task on the following day. We found that, for the learning group, pre-observation (day 1) resting-state FC between bilateral PMd, bilateral M1, bilateral S1 and left SPL was reliably correlated with behavioral scores of motor learning by observing acquired on day 2. Those subjects in the learning group who exhibited greater resting-state FC on day 1 achieved greater motor learning by observing scores on day 2. No such correlation between pre-observation FC and motor learning by observing scores was found for the control group who observed a tutor performing reaches but not learning. Individual differences in grey matter volume could not predict subsequent motor learning by observing behavioral scores. Although the analyses presented here are correlational, the temporal order of events in the experimental design, namely the resting-state scans preceding the observational motor learning task by 24 hours, supports the idea that greater FC in a network linking S1, PMd, M1 and SPL predisposes individuals to learn more about a novel motor skill through visual observation.

The finding that pre-observation resting-state FC between S1 and PMd, M1, and SPL predicts subsequent motor learning by observing is consistent with previous work demonstrating that M1 and the somatosensory system play necessary roles in motor learning by observing. Brown et al. (2009) used repetitive transcranial magnetic stimulation (rTMS) to reduce cortical excitability in M1 immediately after subjects observed a FF learning video. A subsequent behavioral assessment showed that reducing M1 excitability following observation disrupted motor learning by observing. rTMS applied to M1 after observation of FF learning reduced the beneficial effect of observing congruent forces, and eliminated the detrimental effect of observing incongruent forces. These results suggest that M1 plays a key role in motor learning by observing.

We have also recently demonstrated that the somatosensory system plays a necessary role in motor learning by observing (McGregor et al., 2016). We used median nerve stimulation to occupy the somatosensory system with unrelated afferent inputs while subjects observed a video of a tutor undergoing FF learning. During observation, subjects received median nerve stimulation either to the right arm (the same arm used by the tutor in the video), to the left arm (opposite the arm used by the tutor) or no stimulation. Stimulation disrupted motor learning by observing in a limb-specific manner such that stimulation of the right arm (observed effector) interfered with learning, whereas stimulation applied to the opposite arm did not. This result demonstrated that the somatosensory representation of the observed effector is necessary and therefore must be unoccupied during observation for motor learning by observing to occur. In a follow-up EEG experiment, we showed that S1 cortical activity, as assessed using somatosensory evoked potentials, increased for subjects who observed learning by an amount that positively correlated with subsequent behavioral motor learning by observing scores. These results suggest that observation-induced functional changes in S1 support motor learning by observing (McGregor et al., 2016).

The network identified in the current study overlaps with those identified in neuroimaging studies showing that sensory-motor networks support observational learning. We have previously shown that observing motor learning results in changes in resting-state FC between M1, S1, visual area V5/MT and the cerebellum. Functional connectivity changes within this network were correlated with behavioral measures of motor learning, assessed following the fMRI sessions (McGregor and Gribble, 2015). Cross et al. (2009) showed that observation of learned dance movement sequences recruits brain areas including premotor and parietal cortices. The authors reported greater activation in premotor and parietal regions when subjects observed movement sequences on which they had been trained (by observation) over the previous 5 days, compared to untrained movement sequences. These studies suggest that the neural substrates of motor learning by observing include premotor cortex, M1, S1 and parietal cortex. This is consistent with the results of the current study in which subjects who exhibited greater pre-observation resting-state FC between S1 and PMd, M1, and SPL later showed the greatest observation-related facilitation of motor learning.

More generally, the current study provides insight into the neural basis of motor learning. The network identified here closely corresponds to functional networks involved in active motor learning. For example, resting-state fMRI studies of active motor learning have found FC changes between M1, dorsal premotor cortex and the cerebellum following FF adaptation (Vahdat et al., 2011) and FC changes within the fronto-parietal resting-state network following visuomotor adaptation (Albert et al., 2009). Several task-based neuroimaging studies have similarly suggested a role for PMd (e.g., Steele and Penhune 2010), M1 (e.g., Grafton et al., 1992; Steele and Penhune 2010), S1, and SPL in motor learning through active movement (see Hardwick et al., 2013 for review).

There are commonalities between the functional network identified in the current study and those functional networks that have been previously reported to predict aspects of motor learning through active movement training. Tomassini et al. (2011) showed that the task-based activation of premotor and parietal cortices (along with prefrontal cortex, basal ganglia and the cerebellum) is associated with higher behavioral measures of motor learning. Wu et al. (2014) have similarly shown that resting-state FC (as measured by high-density EEG) between M1, premotor cortex and parietal cortex can predict skill acquisition. The consistency between predictive functional networks for learning through active movement training and observational motor learning provides evidence in favor of similar neural substrates for these two forms of motor learning.

There is evidence from the motor learning literature that individual differences in brain structure can predict learning through active practice. Tomassini et al. (2011) demonstrated that individual differences in grey matter volume within the cerebellum and higher order visual areas (V2, V3, V5/MT) can also predict behavioral measures of motor learning during a visuomotor tracking task. While there is evidence for structure-based predictability of active motor learning, in the current study we found that this was not the case for motor learning by observing; individual differences in grey matter volume could not account for variability in behavioral scores of motor learning by observing. The discrepancy between the results of the current study and that of Tomassini et al. (2011) may be due to methodological differences in terms of the T1-image acquisition parameters and VBM analysis procedures used, and/or the current study may have had insufficient statistical power. Future studies investigating grey matter volume correlates of motor learning by observing should have a larger sample size to increase statistical power.

Here we tested if pre-observation measures of brain function or structure could predict subsequent motor learning by observing. We found that pre-observation resting-state FC between bilateral S1, PMd, M1, and left SPL predicted the extent to which observation would promote motor learning on the following day. Individual differences in grey matter volume could not predict behavioral scores of learning following observation. These results demonstrate that individual differences in resting-state FC among sensory-motor cortical brain areas can explain part of the individual variability in the extent to which observation facilitates motor learning. This finding is consistent with the idea that those individuals who have more 'primed' sensory-motor circuits are more predisposed to motor learning through observation. Pre-observation FC within the identified sensory-motor network may be used as a biomarker of the extent to which observation will promote motor learning. Predicting an individual's predisposition for motor learning by observing could be valuable in a clinical context for planning individualized rehabilitation strategies and improving prognostic accuracy (Stinear, 2010).

The origin of individual variability in pre-observation sensory-motor FC is still unclear. In one scenario, it is possible that the observed individual differences in FC are a reflection of functional variability and not anatomical variability within this network. However, given the close correspondence between anatomical and functional connectivity (e.g., Fox et al., 2005), another scenario is that the observed differences in FC arise from individual differences in anatomical connectivity. For example, it could be the case that greater structural connectivity between these sensory-motor brain areas results in higher pre-observation sensory-motor FC which, in turn, promotes greater motor learning by observing. Since we did not acquire images for performing structural connectivity-based analyses (such as diffusion tensor images) in the current study, we cannot rule out the possibility that individual differences in structural connectivity among sensory-motor brain areas underlies the effect seen here, whereby pre-observation FC predicts motor learning by observing.

However, resting-state FC does not only reflect anatomical connectivity. Indeed, much work has shown that resting-state FC can be shaped by recent experiences. Such "stimulus-rest interactions" have been demonstrated across several domains. For example, exposure to visual stimuli (Lewis et al., 2009) or undergoing active motor learning (Albert et al., 2009) can change resting-state FC. Since resting-state FC is affected by both structure and function, it is likely the case that both of these factors contribute to individual differences in pre-observation sensory-motor FC. While we cannot further pursue this question using the current dataset, this would be an interesting avenue for future research. Since previous experiences can alter resting-state FC, it is likely that performance of null field reaches in the baseline condition ‘primed’ sensory-motor networks prior to the day 1 resting-state scans and perhaps increased the sensitivity of the current study. It would be of interest to examine if baseline measures (i.e., without prior null field reaches) of resting-state FC within the identified could also predict motor learning by observing. Another outstanding issue is the stability of these individual differences in pre-observation FC over time. Future research should examine the test-retest reliability of pre-observation FC over longer time periods (e.g., several days or weeks apart) to establish the long-term stability of the FC patterns within the network presented here. This would allow one to better distinguish between within-session patterns from those more permanent structural or functional patterns.

## Acknowledgements

This work was supported by the Natural Sciences and Engineering Research Council of Canada, and by the National Institute of Child Health and Human Development R01 HD075740

